# The high infectivity of SARS-CoV-2 B.1.1.7 is associated with increased interaction force between Spike-ACE2 caused by the viral N501Y mutation

**DOI:** 10.1101/2020.12.29.424708

**Authors:** Jadson C. Santos, Geraldo A. Passos

## Abstract

The Spike glycoprotein receptor-binding domain (RBD) of SARS-CoV-2 mediates the viral particle’s binding to the angiotensin-converting enzyme 2 (ACE2) receptor on the surface of human cells. Therefore, Spike-ACE2 interaction is a crucial determining factor for viral infectivity. A new phylogenetic group of SARS-CoV-2 (lineage B.1.1.7) has been recently identified in the COVID-19 Genomics UK Consortium dataset, which features an amino acid substitution in the Spike RBD (N501Y mutation). Infections with the SARS-CoV-2 lineage B.1.1.7 have been overgrowing in recent weeks in the United Kingdom, indicating an even greater spread capacity than that seen with previous strains of the novel coronavirus. We hypothesized that this rapid spreading/infectivity of the B.1.1.7 lineage might be due to changes in the interaction force between the mutant Spike RBD and ACE2. This study employed *in silico* methods involving mutagenesis (N501Y mutation) and interface analysis focusing on the Spike RDB-ACE2 interaction. The results showed that the SARS-CoV-2 N501Y mutant (lineage B.1.1.7) establishes a more significant number of interactions relating to the mutant residue Y501 (Spike RDB) with residues Y41 and K353 (ACE2). This finding shows that the increased infectivity of SARS-CoV-2 lineage B.1.1.7 is associated with the interaction force between the Spike RBD Y501 mutant residue with the ACE2 receptor, which in this strain is increased.

## Introduction

The fast SARS-CoV-2 spreading among humans is pushing its molecular evolution. So far, the virus has accumulated mutations at a rate of up-to-two-nucleotide changes every month. Recent isolates of the SARS-CoV-2 feature at least 20 nucleotide changes in their genomes than the earliest isolates in January 2020 (Rambaut et al., 2020).

Most of the variability found in SARS-CoV-2 isolates feature mutations within the Spike receptor-binding domain (RBD), a portion of the Spike glycoprotein that mediates viral attachment to the angiotensin-converting enzyme 2 (ACE2) receptor on the surface of human cells (Shang et al., 2020).

The emergence of the SARS-CoV-2, B.1.1.7 lineage, detected in early December 2020 by the COVID-19 Genomics UK Consortium (https://www.cogconsortium.uk/) (Kupferschmidt, 2020; Rambaut et al., 2020;) represents an example, among several other, of the fast viral molecular evolution.

The B.1.1.7 lineage is somewhat surprising, as it accumulates 17 mutations in its genome. Eight of them are located in the gene that encodes the Spike protein on the virus surface (Rambaut et al., 2020). The RBD is crucial for Spike docking with the ACE2 receptor on the surface of human cells (Kemp et al., 2020). B.1.1.7 lineage phenotype has also attracted attention, as it proves to be much more transmissible among humans than the other known SARS-CoV-2 lineages (Kupferschmidt, 2020; Rambaut et al., 2020;).

The N501 has been identified as a residue that increases the binding affinity to human ACE2 (Starr et al., 2020). According to Rambaut et al. (2020), the mutation N501Y found in the B.1.1.7 lineage, which is located at the Spike gene, covers one of the six key contact amino acid residues within the Spike RBD.

Determinations of the Spike RDB domain’s surface affinity strength with the ACE2 receptor have enabled mapping of the amino acid residues that may represent desired targets for the development of vaccines against COVID-19 or antibody-based therapies against SARS-CoV-2 (Sternberg and Naujokat, 2020).

However, quantitative impact of the SARS-CoV-2 N501Y mutation on the Spike RBD affinity to the ACE2 receptor is still unknown.

In this study, we raised the hypothesis that N501Y mutation increases the interaction force between Spike RBD and ACE2 receptor.

To test this, we used *in silico* tools to mutagenize and cause the N501Y mutation in the Spike RDB domain, at the protein level, and simulate the molecular interactions with the ACE2 receptor. Comparisons between wild type versus mutant Spike RDB structures showed that the mutant N501Y RDB interacts with greater affinity to the ACE2 receptor.

## Methods

### Spike RDB-ACE2 interface analysis

This study uses the structure deposited in the Protein Data Bank available at (https://www.rcsb.org/), code 6M0J. The PyMOL software, available at (https://pymol.org/2/), was used to: 1) visualize the interaction between the amino acid residue N501 of the Spike RDB domain (SARS-CoV-2, isolate Wuhan-Hu-1, NCBI reference genome sequence NC_045512.2) and the Y41 residue of the human ACE2 receptor protein, 2) simulate the N501Y mutation found in the isolate SARS-CoV-2 lineage B.1.1.7 (Rambaut et al., 2020) and 3) analyze the interactions resulting from the Spike (RDB) N501Y mutation.

Moreover, we used PDBePISA software (Krissinel and Henrick, 2007), available at (https://www.ebi.ac.uk/pdbe/pisa/) for a detailed analysis of the Spike RDB-ACE2 quaternary structures, comparing the interaction resulting from the Spike N501 or Y501 mutant residue regarding their chemical properties and the predicted dissociation pattern.

### Statistical analyses

The server PDBSum (Laskowski et al., 2018), available at (http://www.ebi.ac.uk/thornton-srv/databases/pdbsum/Generate.html), was used for comparative statistics of the interfaces formed between the Spike N501, or the respective N501Y mutant residue, with ACE2 receptor.

## Results

### Spike RBD-ACE2 interface visualization

The original SARS-CoV-2 Spike RBD-ACE2 interface region feature the N501 amino acid residue (Spike RBD) that establishes two non-bonded contacts with the Y41 and K353 residues (ACE2) at 3.5-angstrom distance as predicted by the PyMOL and PDBsum softwares (Figure 1).

**Figure 1.**
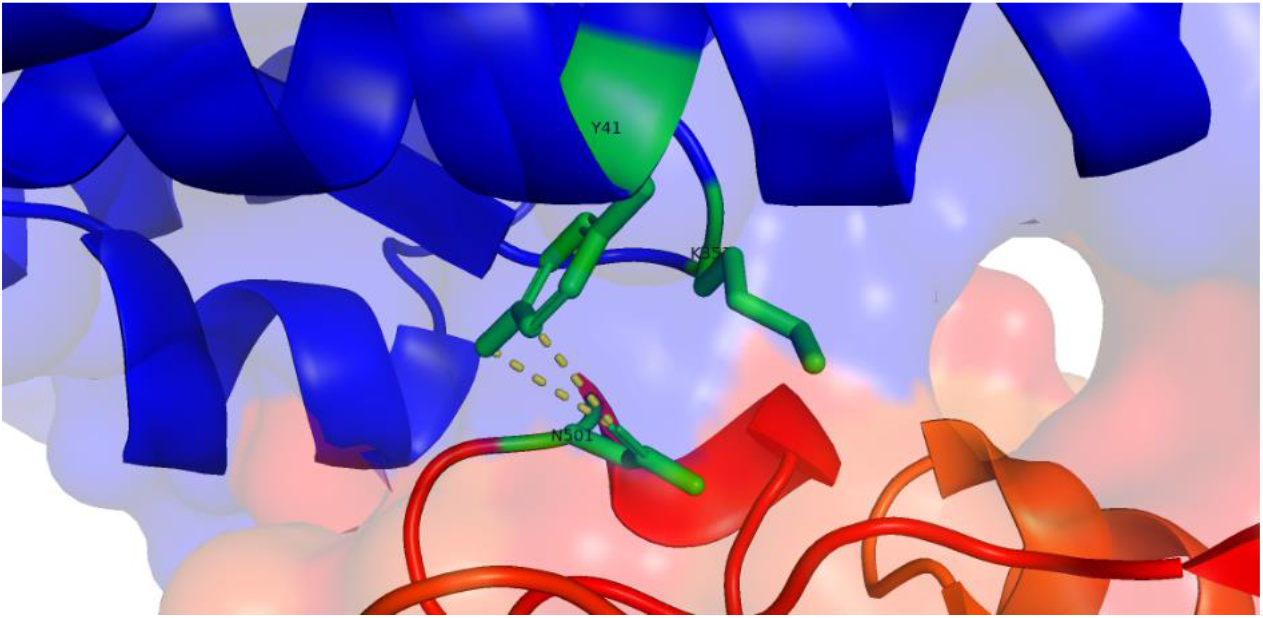
SARS-CoV-2, Wuhan-Hu-1 lineage, Spike RBD-ACE2 interaction interface featuring the N501 amino acid residue (on the Spike RBD side in red) that establishes two non-bonded interactions with Y41 residue (on the ACE2 side in blue) at 3.5 angstrom distance. Predictions calculated with PyMOL.

In contrast, PyMOL and PDBsum indicates that the mutant SARS-CoV-2 N501Y (lineage B.1.1.7) features the Y501 residue (Spike RBD) that establishes one hydrogen bond with K353 residue and non-bonded interactions with Y41, D38 and K353 residues of the ACE2 receptor at 3.5-angstrom distance (Figure 2).

**Figure 2.**
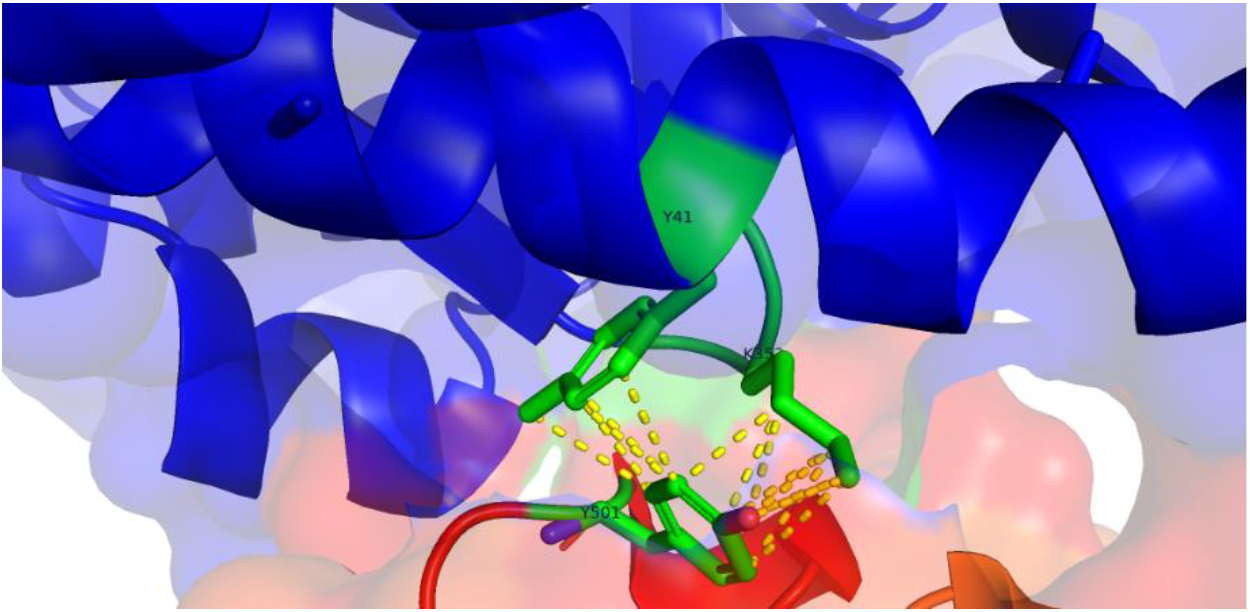
SARS-CoV-2, mutant N501Y, lineage B.1.1.7, Spike RBD-ACE2 interaction interface featuring the Y501 mutant amino acid residue (from the Spike RBD side in red) that establishes one hydrogen bond with K353 residue and non-bonded interactions with Y41 and K353 residues from the ACE2 side (in blue), at 3.5 distance. Predictions calculated with PyMOL.

In addition, PDBePISA analysis predicted the interactions at the interface amino acid residue N501 Spike(RBD)-ACE2 (Figure 3).

**Figure 3.**
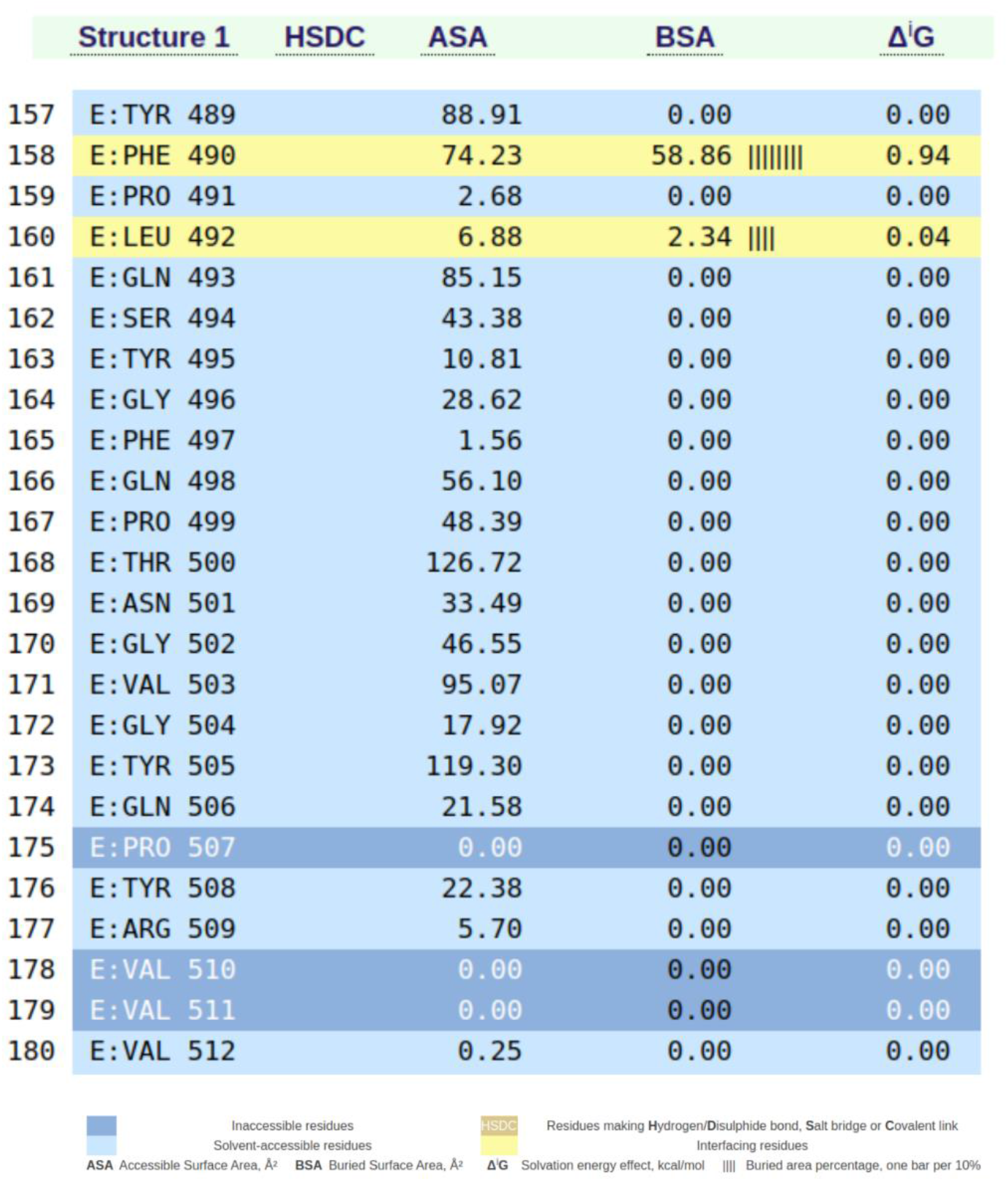
PDBePISA prediction analysis showing part of the interactions at the N501 residue Spike(RBD)-ACE2 interface.

Moreover, increased interactions occurred at the N501Y mutant residue Spike(RBD)-ACE2 interface. Were identified supplementary hydrogen bonds involving the residues Y500, Y501, G502, V503, and Y505 from the Spike RBD domain side, which increase the interaction force with ACE2 receptor and the RBD domain, buried surface area (BSA) (Figure 4).

**Figure 4.**
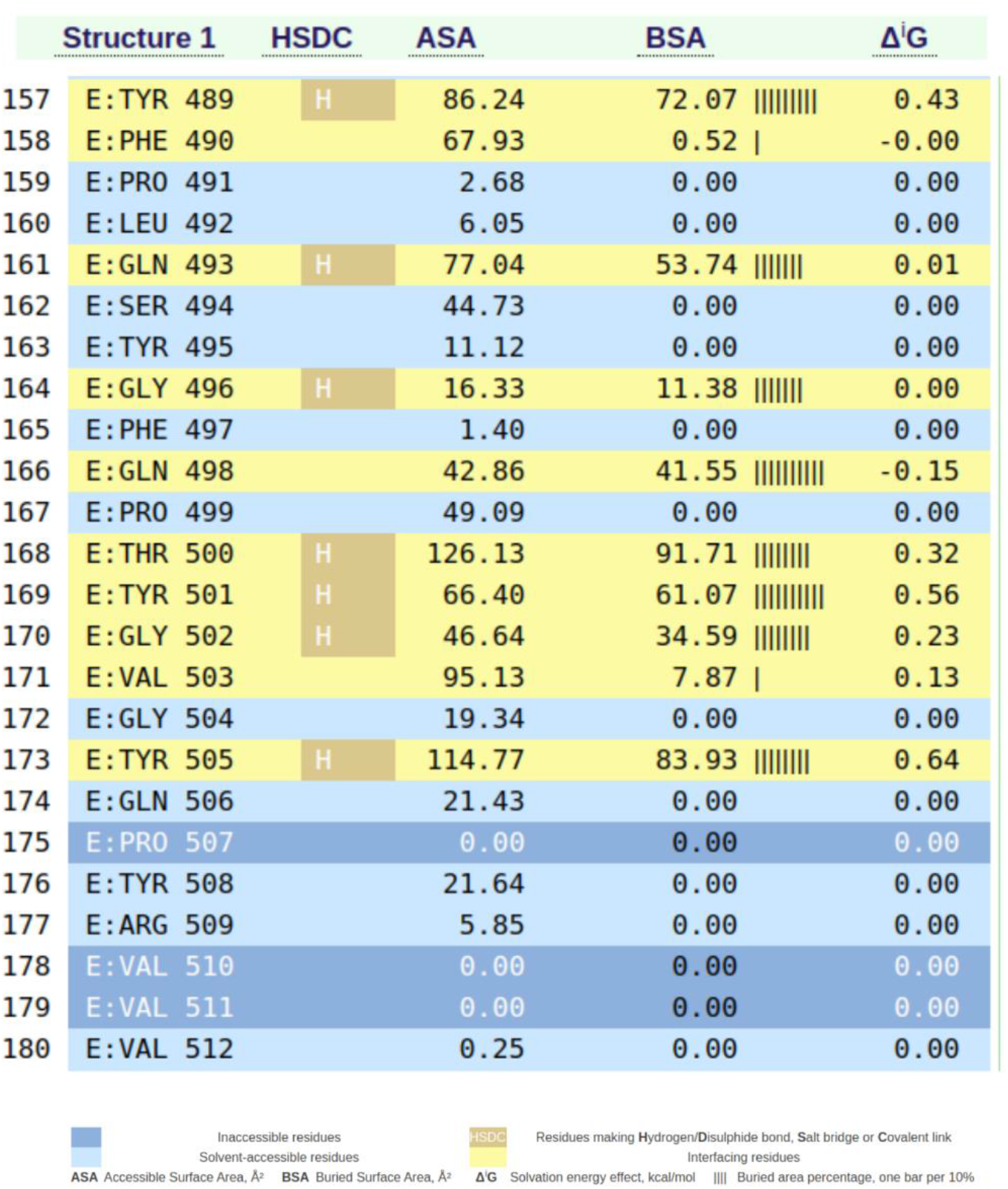
PDBePISA prediction analysis showing the possibility for increased interactions at the N501Y mutant Spike(RBD)-ACE2 interface. The surrounding 501Y residues (Y500, G502, V503 and Y505) feature increased interaction force with the ACE2 receptor.

### Statistical analysis of the RBD-ACE2 interface

The chemical nature of the interactions between amino acid residues was predominantly non-covalent, as determined by the PDBSum software. The mutant Spike RBD Y501 residue resulted in greater apolar and hydrogen bonds with ACE2 Y41 and K353 residues. Noteworthy, the SARS-CoV-2 B.1.1.7 lineage establishes a more significant number of interactions with the ACE2 receptor. In addition to the mutant Y501 residue, the non-mutant border residues, such as Y500, G502, V503, and Y505, show polar interactions with the ACE2 receptor (Figure 5).

**Figure 5.**
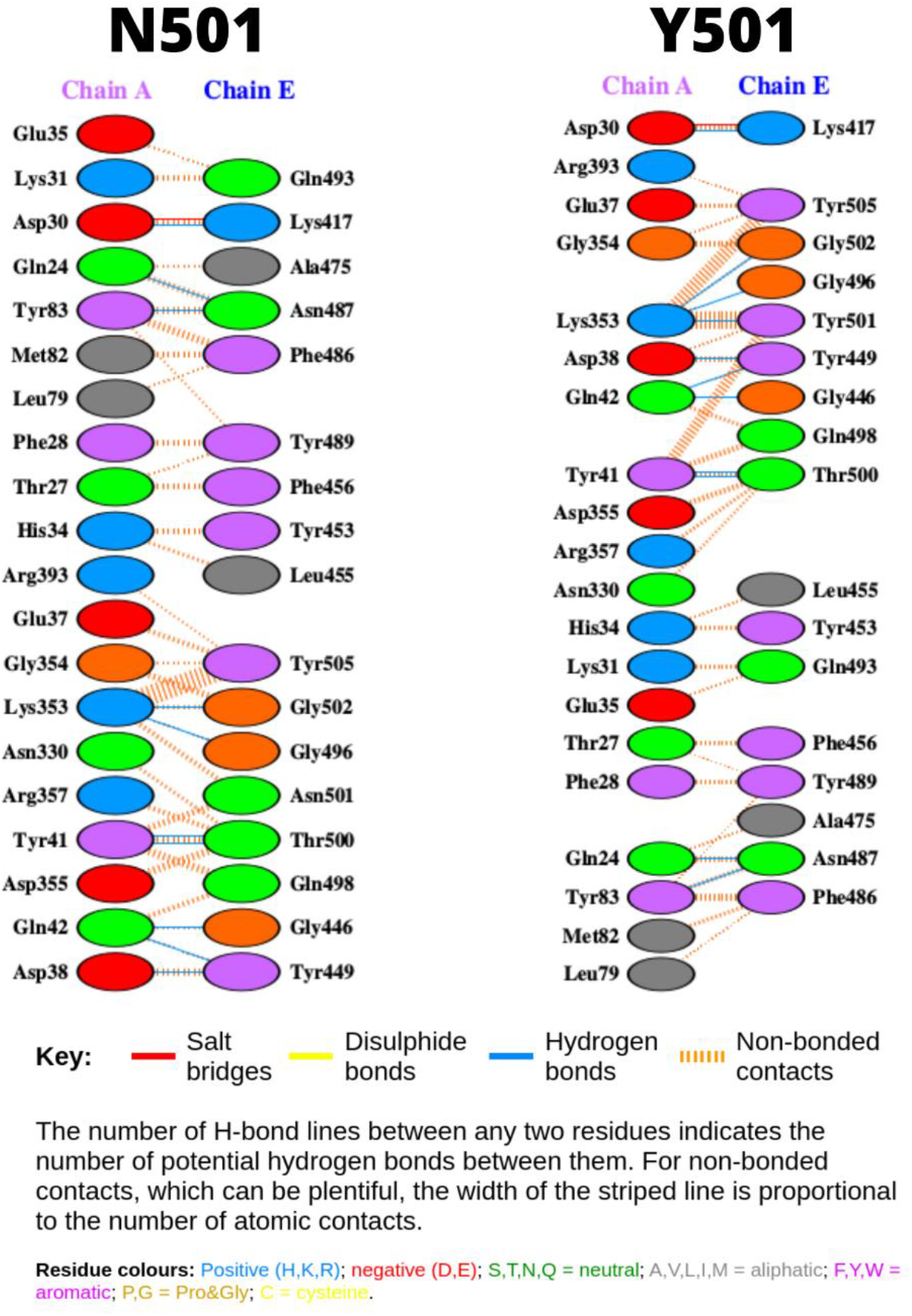
Number and chemical nature of the interactions between amino acid residues N501 (Wuhan-Hu-1 lineage) or Y501 (B.1.1.7 lineage) with ACE2 receptor. Most of the interactions were non-covalent, as determined with the PDBSum software. (Chain A: ACE2 receptor, Chain E: Spike RBD).

## Discussion

This study shows through *in silico* protein-protein interface analysis that the mutation N501Y in the Spike receptor-binding domain (RBD) of the SARS-CoV-2, lineage B.1.1.7, establishes increased interaction with the ACE2 receptor when compared to the Wuhan-Hu-1 lineage.

The lineage B.1.1.7 is highly transmissible and infectious among humans (Rambaut et al., 2020). Since the beginning of December 2020, its spreading in the United Kingdom significantly increased (Rambaut et al., 2020; Kupferschmidt, 2020).

The N501Y mutation occurs in the receptor-binding domain (RDB) of the Spike protein of the SARS-CoV-2, a crucial region during the virus’s interaction with the ACE2 receptor on the surface of the human cells and progress with the infection (Starr et al., 2020).

Therefore, we hypothesized that the mutation N501Y, found in the strain B.1.1.7, could cause the Spike protein to establish stronger interactions with the ACE2 receptor.

The hypothesis was confirmed when we performed the Spike (RBD)-ACE2 interaction analysis by comparing SARS-CoV-2 (lineage Wuhan-Hu-1) Spike RBD residue N501 versus the mutant residue Y501, lineage B.1.1.7.

The nature of the interactions was predominantly non-covalent and involving hydrogen bonds, polar and apolar bonds.

Interestingly, the N501Y mutation causes a change in the spacing between the amino acid residues of the Spike protein, represented by the Buried Surface Area in PDBePISA. We observed that this type of alteration causes the amino acid residues around the 501Y residue to establish more interactions with the ACE2 receptor.

Together, these changes cause the Spike protein in the RBD segment of strain B.1.1.7 to interact more strongly with the ACE2 receptor.

The greater Spike-ACE2 interaction strength may support a better understanding of the high infectivity and spread the SARS-CoV-2, lineage B.1.1.7, or other strains featuring similar characteristics, based on the molecular affinity between the virus and its host cells.

## Acknowledgments

This study was funded in part by the following agencies; Fundação de Amparo à Pesquisa do Estado de São Paulo (FAPESP, São Paulo, Brazil) through grant No. 17/10780-4, Conselho Nacional de Desenvolvimento Científico e Tecnológico (CNPq, Brasília, Brazil) through grant No. 305787/2017-9 and Coordenação de Aperfeiçoamento de Pessoal de Nível Superior (CAPES, Brasília, Brazil) through financial code 001. JCS received a fellowship from CAPES.

## Author’s contributions

GAP conceived and supervised the project and wrote the manuscript. JCS selected the raw sequence data and bioinformatics tools, designed and used the pipeline, and analyzed the data.

